# Single-cell analyses reveal a simple multi-gene transcriptomic signature with predictive power in prognosis and therapy effectiveness in triple-negative breast cancer

**DOI:** 10.64898/2026.08.12.744411

**Authors:** Guillaume Davidson, Véronique Debien, Tom Sexton

## Abstract

Triple-negative breast cancer (TNBC) is an aggressive, heterogeneous form of breast cancer with limited specific therapy options, prevalent metastasis and frequent relapse. Re-analysis of single-cell RNA-sequencing data characterizes the diverse cell subtypes within the tumor and microenvironment of TNBC, supporting a luminal progenitor origin for the cancer and providing clues as to the factors involved in progression of the disease. The relative burdens of these subtypes can be deconvolved from bulk RNA-sequencing data, readily identifying the stem-like, mesenchymal and stromal cell subtypes significantly associated with poor survival and enrichment in metastasis. Importantly, these can be simplified to ten-gene signatures with comparable predictive power, notably in response to different therapeutic strategies, which are linked to relative burdens of different subtypes of stromal fibroblasts. The expression level of these signatures could provide a cheap means for selecting therapy strategies in personalized medicine.

## BACKGROUND

Breast cancer is one of the most frequent cancers in women and its incidence rate has been steadily increasing [1]. Three main subtypes are defined based on protein expression assessed by immunohistochemistry results: hormone receptor positive (estrogen and/or progesterone receptor positive (ER+/PR+) human epidermal growth factor receptor 2 (HER2)-negative, referred to as luminal breast cancer, HER2-positive and triple-negative breast cancer (ER-, PR-, HER2-; TNBC) [2]. TNBC is an aggressive subtype with high metastatic capacity and is prone to relapse [3]. Metastasis and treatment resistance mechanisms have been studied in all cancer types and frequently rely on plasticity of cancer cells and interplay with the tumor microenvironment (TME) [4]. In TNBC, the presence of intratumoral cancer stem cells (CSCs) has been linked to disease progression. These cells can appear through epithelial-to-mesenchymal transition (EMT) of cancer cells and display different phenotypes identified through markers such as CD44+/CD24-or ALDH+ [5]. The complex CSC-TME interactions can affect tumor progression and influence how the tumor responds to different therapies. [6]. Thus, comprehensive profiling of cancer cell states and their associated TME helps characterize tumoral heterogeneity, which may better inform personalized therapeutic options. In routine practice, due to lack of HER2 or hormone receptor expression, biomarker-driven treatment options are more limited for TNBC than for other subtypes. Chemotherapy remains the therapeutic backbone despite ongoing evaluation of new strategies [7], and recent immunotherapy-containing approaches have become standard of care for a selected proportion of patients with metastatic and early TNBC [8]. However, despite these recent advances, treatment resistance and relapse are still major problems, and TNBC classifications based on bulk transcriptomics alone have so far not been widely adopted to inform clinical decisions [9–11]. In this study, we aimed to derive simplified transcriptomic signatures from single-cell RNA-sequencing data that capture clinically relevant malignant and stromal cell states in TNBC and can be assessed in more readily available bulk transcriptomic datasets. To this end, we re-analyzed publicly available single-cell RNA-sequencing data from over 50,000 cells of TNBC samples to characterize the tumor cell types and cellular populations within the TME. We subsequently reduced the identified cell-state specific expression programs to selected gene signatures and evaluated their associations with patients’ outcomes and response to neoadjuvant chemo or chemo-immunotherapy in independent transcriptomic cohorts. Through this approach, we sought to bridge the biological resolution of single-cell profiling with a potentially more scalable framework for prognostic and treatment-response stratification in TNBC.

## METHODS

### Processing and analysis of single-cell RNA-sequencing data

The eight TNBC samples (patients 0106, 0135, 0126, 0114, 4031, 0131, 0554, 0177) and two normal tissue samples (patients 0123, 0093) were downloaded from GEO (GSE161529) [12]. For each sample, barcodes, features and matrix cell ranger outputs were processed individually in R v4.4.3 using Seurat v5.3.0 [13] to filter out poor quality cells. Cells were filtered to keep only those with feature counts ranging from 200-5000 and a proportion of mitochondrial reads < 20%. Potential doublets were removed using package DoubletFinder [14], assuming a doublet rate of 0.8% per 1000 cells reported by cell ranger. After filtering, samples were merged (“merge()”), counts were log-normalized (“NormalizeData()”), variable features were determined (“FindVariableFeatures()”), and data were scaled (“ScaleData()”) before dimension reduction (“RunPCA()”). In order to mitigate batch effects, the Harmony method [15] implemented in Seurat5 was used during the integration step by providing the “HarmonyIntegration” argument to “IntegrateLayers()”, then clustering was performed to a resolution of 0.8 (“FindNeigbors()” and “FindClusters()”) and UMAP (uniform manifold approximation and projection) projection was computed (“RunUMAP()”) with the 40 most significant components. Layers were merged (“JoinLayers()”) and cluster marker genes were calculated (“FindAllMarkers()”), then filtered (log2(fold-change) > 1; adjusted p-value < 0.05; pct1 > 20%). For each broad cell type (lymphoid, myeloid, fibroblastic, epithelial and cancer), re-clustering was performed using the same procedure after selecting only relevant cells. In re-clustering analyses, some small clusters mainly showed high expression of *MALAT1* and *XIST*, these were deemed poor quality cells and filtered out. To represent gene expression on the global UMAP without overlapping signal, the schex page (https://github.com/SaskiaFreytag/schex) was used. To represent gene expression on UMAPs from re-clustering analyses, Seurat function “FeaturePlot()” was used. Gene regulatory networks and transcription factor activity were determined using SCENIC+ [16] with python v3.7 and processed in R using SeuratExtend v1.2.1 [17]. The final cluster lists can be found in **Table S1** and their corresponding marker genes in **Table S2**.

### Processing and analysis of bulk RNA-sequencing data

Raw RNA-seq read counts from the TCGA-BRCA collection [18], Garcia-Recio *et al.* [19], and the SCANDARE (NCT03017573) TNBC cohort [20] were downloaded from the GDC data portal, GEO (GSE209998) and EGA (EGAS50000000970), respectively. In all cohorts, only samples labeled as TNBC with full clinical annotations were kept (n = 1069; 129 and 114 samples; n = 177, 52 and 114 TNBC). The raw-count matrices were normalized by sequencing depth by the median-of-ratios method implemented in the DESeq2 package v1.42.1, and then normalized counts were divided by median transcript length, based on genome assembly hg38 with ENSEMBL v101 annotations. In the SCANDARE study, resistant or sensitive samples are assessed by the response to treatment at surgery by the residual cancer burden (RCB) index: Patients with an RCB index of 0 or I were categorized as chemo-sensitive and those with an RCB index of II to III were defined as chemo-resistant.

### Analysis of microarray data

Agilent 44K microarray normalized gene expression data from the I-SPY2 clinical trial [21] were downloaded from GEO (GSE194040). Samples were filtered to keep only TNBC from the neoadjuvant Paclitaxel + Pembrolizumab treatment arm (n = 987 samples, n = 362 TNBC, n = 29 Pembrolizumab arm).

### Functional analysis of sequencing data

Deconvolution from single-cell data was performed using the web-tool version of CIBERSORTx [22] with absolute quantification and S-mode. In order to reduce complexity of the task, close clusters were merged together: TNBC.LPlike1 and TNBC.LPlike2 to “TNBC.LPlike”, TNBC.MESlike and TNBC.IFN to “TNBC.MESlike-IFN”, and TNBC.hypoxia and TNBC.inf to “TNBC.hypoxia-inf”. Normal breast tissue clusters with high expression of hormone receptors (“Mature Lumen” and “Luminal Progenitor type 3”) tended to show high bias in some samples, in particular for the Luminal Androgen Receptor (LAR) subtype, so these were excluded. Additionally, clusters with no unique markers such as clusters engaged in cell cycle and intermediate fibroblasts were also excluded (TNBC.cycling, Myeloid.cycling, Lymphoid.cycling, FIB.int). To build gene signatures for each relevant cluster, the top ten genes were selected based on adjusted p-value computed by Seurat and keeping only one gene for each family. Signature values were computed in bulk sequencing data by using the geometric mean of each list. Sequencing data were represented as heatmaps with pheatmap package v.1.0.13, and as bar plots and box-and-whisker plots with the ggplot2 package v4.0.2. Gene ontology analyses were performed using the web-tool DAVID [23].

### Survival analysis

Relationships between CIBERSORTx score or gene signatures and patient survival (overall survival, OS) were computed in Rv4.5.2 using packages survival v3.8.3 and survminer v.0.5.2. Patients were stratified based on CIBERSORTx scores or signature values using “surv_cutpoint()” and Kaplan-Meier curves were represented with the “survfit()” and “ggsurvplot” functions. Hazard ratios were determined by univariate Cox proportional hazards model with the “coxph” function.

## RESULTS

### Diversity of cell types within TNBC tumor and microenvironment

To characterize the different cell types present in TNBC, we re-analyzed a single-cell transcriptomic study of a human cohort [12], focusing on the eight TNBC tumor samples and two normal tissue samples, corresponding to a total of 56,696 cells after quality filtering. Cells clustered into five broad types: epithelial, endothelial, fibroblast, myeloid and lymphoid cells, supported by the expression of marker genes within specific clusters (**Fig 1a,b; Fig S1a**). These cell types were present across all tumor samples; the normal breast tissue was predominantly epithelial (**Fig S1b,c**). Re-clustering the epithelial (*EPCAM*+) cells revealed populations of normal breast epithelial cells, including basal cells (“Basal”; *DKK3*+), mature luminal cells (“ML”; *FOXA1*+) and three clusters of luminal progenitor cells (“LP1”, “LP2”, “LP3”; *SOX10*+); the 23,145 epithelial TNBC cells re-clustered into eight groups with distinct phenotypes (**Fig 1c**; **Fig S1d,e; Fig S2; Tables S1, S2**). Two of these shared marker genes with luminal progenitors, most closely resembling the LP2 cluster (“TNBC.LPlike1”, “TNBC.LPlike2”; *SOX10*+, *EHF*+). The remaining TNBC-specific clusters could be characterized as a stem-like phenotype (“TNBC.STEMlike”; *ALDH1A1*+), a mesenchymal-like state (“TNBC.MESlike”; *FN1*+), an interferon-response state (“TNBC.IFN”; *IFI6*+, *IFI44*+, *STAT1*+), an inflammation-response state (“TNBC.inf”; *CD74*+), a hypoxia-response state (“TNBC.hypoxia”; *VEGFA*+), and cycling cells (“TNBC.cycling”; *MKI67*+). Looking closer at expression of key markers across clusters, it is apparent that the four clusters TNBC.MESlike, TNBC.IFN, TNBC.inf and TNBC.hypoxia are fairly dedifferentiated and express hypoxic and inflammatory response programs (**Fig 1d**). Additionally, TNBC.MESlike is very close to TNBC.IFN and also expresses interferon-response genes. These four proximal clusters are reminiscent of previously described expression modules which have been associated with cancer recurrence in many different tumor types [24]. Despite their different phenotypes, we find some genes that are expressed across all cancer-specific clusters, including *FOLR1*, found to be associated with growth of TNBC cell lines [25], and *ST3GAL4*, linked to aerobic glycolysis *in vitro* [26].

**Figure 1.**
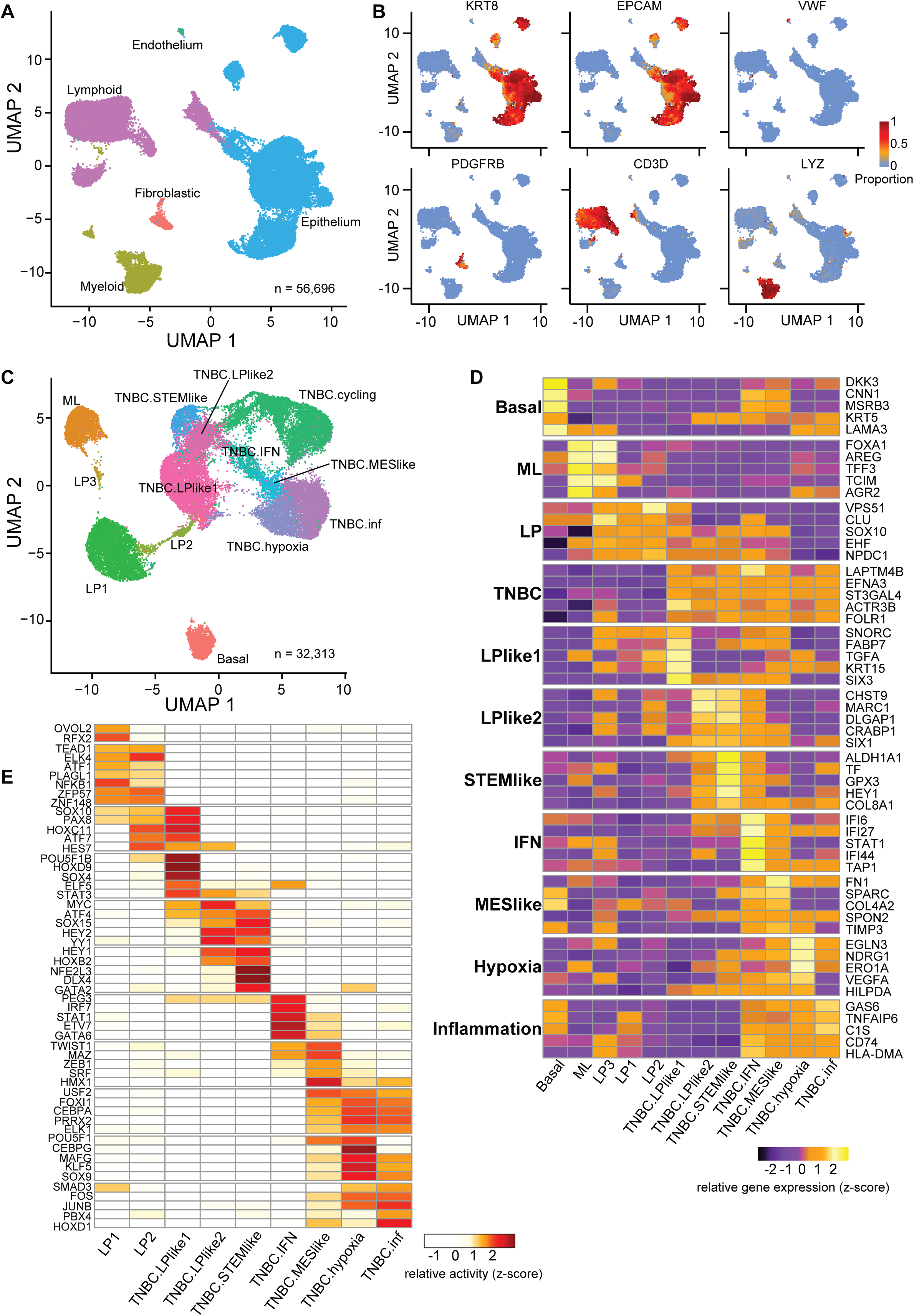
Cell diversity within tumor and microenvironment of TNBC. **a**) UMAP projections and clustering of single-cell transcriptomic data (from ref 12) comprising eight TNBC and two normal samples. **b**) The same UMAP as **a**), with indications of the proportions of cells expressing the denoted marker genes. **c**) UMAP projections and re-clustering of the EPCAM^+^ (epithelial) portion of the cells from **a**). **d**) Heat map showing the relative expression levels of given marker genes within the different epithelial clusters, identified in **c**). **e**) Heat map showing the relative activity of denoted transcription factors in the gene regulatory networks for the epithelial clusters, identified in **c**), after SCENIC analysis.

Of the four non-epithelial cell groups (predominantly found in tumor samples), three could be re-clustered into distinct subtypes; endothelial cells were not numerous and comprised a single cluster (“ED”; *VWF*+, *EMCN*+). Myeloid cells separated into eleven clusters (**Fig S3a,b**), including classical monocytes (“Mono.cla”; *VCAN*+, *CD16*-), intermediate monocytes (“Mono.int”; *S100A8*-high), atypical monocytes (“Mono.aty”; *CD16*+), tumor-associated macrophages (“TAM”; *MRC1*-high), conventional dendritic cells (“DC.CD1C”, *CD1C*-high), interleukin-12-producing dendritic cells (“DC.CD1A”, *CD1A*-high), immunomodulatory dendritic cells (“DC.LAMP3”, *LAMP3*+), plasmacytoid dendritic cells (“DC.pla”, *LILRA4*+), granulocytes (“Granulocyte”, *TACSTD2*+), mast cells (“Mast”; *CPA3*+), and cycling cells (“Myeloid.cycling”; *MKI67*+). Atypical monocytes and TAMs displayed an M2 macrophage signature and high expression of *TREM2*, previously associated with immunosuppression promoting tumor progression [27]. Lymphoid cells separated into thirteen clusters (**Fig S3c,d**), including two memory/naïve CD4+ T cell clusters (“TCD4.IL7R”, “TCD4.LTB”, *IL7R*+, *LTB*+, respectively), tertiary lymphoid structure-associated CD4+ T cells (“TCD4.CXCL13”; *CXCL13*+), regulatory CD4+ T cells (“TCD4.reg”; *FOXP3*+), cytotoxic effector CD8+ T cells (“TCD8.eff”; *GZMK*-high), natural killer-like CD8+ T cells (“TCD8.NKT”; *CD8A*+, *KLRD1*+), natural killer cells (“NK”; *KLRD1*+), follicular B cells (“Bcell”; *CD79*+, *MS4A1*+), four plasma cell clusters (“PC.MZ1B”, “PC.IGLC”, “PC.IGHA”, “PC.IGHV”; *CD79*+, *SDC1*+), and cycling cells (“Lymphoid.cycling”; *MKI67*+). The “TCD8.eff” and “TCD8.NKT” clusters displayed high expression of exhaustion markers, such as *LAG3* and *HAVCR2*.

Fibroblast cells separated into five clusters (**Fig S3e,f**), including pericytes (“FIB.peri”; *RGS5*+), fibro-adipogenic progenitors (“FIB.adi”; *APOD*+), myofibroblastic-type cancer-associated fibroblasts (“FIB.myCAF”; *FAP*+), hypoxic CAFs (“FIB.hypoxia”; *VEGFA*+), and an intermediate state in between myCAFs and fibro-adipogenic progenitors (“FIB.int”). The “FIB.adi” cluster displayed the CAF progenitor gene program reported in previous studies [28], and are likely derived from adipocytes that are missing in the dataset.

### A possible ontogeny of TNBC progression from luminal progenitors

Previous single-cell transcriptome studies of breast cancer, using the prior subtype classification [9] as a reference, determined that TNBC predominantly has “basal-like” expression profiles, closest to basal epithelia [29]. However, independent lines of evidence strongly favor luminal progenitors as the cell of origin for TNBC [30–32], which is also mirrored by the higher proximity between TNBC and luminal progenitor clusters in the UMAP (**Fig 1c**). To better understand the possible ontogeny of TNBC progression, we performed differential gene expression analysis between “normal” LPs (LP1), “pre-neoplastic” LPs (LP2) and LP-like TNBC (TNBC.LPlike1) (**Tables S2, S3**). We found 44 genes upregulated in LP2 compared to LP1. These were mainly involved in stress response, and seven encode transcription factors (TFs) such as *JUN*, *FOS*, *ATF3* and *MYC*. When comparing LP2 to LP-like TNBC, 628 genes were upregulated in TNBC, involved in developmental programs like Notch signaling (*NOTCH3*, *JAG1*) and Wnt signaling (*CTNNB1*, *MDK*), and encoding TFs of the AP-1 (*JUNB*, *FOSB*, *FOS*), Sox (*SOX4*, *SOX6*, *SOX8*) and Six (*SIX3*) families. A role for these established drivers of oncogenesis is also likely in the conversion of normal luminal progenitors into carcinoma, whose mistakenly “basal” profile could be attributed to subsequent upregulation of mesenchymal markers.

To identify transcriptional regulators governing transformation and different TNBC phenotypes, we performed SCENIC analysis [16] on cells of the epithelial compartment (**Fig 1e**). We found that RFX2 and the epithelial identity regulator OVOL2 were active in the LP1 cluster, but were lost in preneoplastic LP2 and TNBC clusters. In line with this, OVOL2 has previously been described as critical in maintaining epithelial identity and restraining EMT [33]. As expected for their apparent phenotypes, the interferon-response TNBC cluster showed activation of interferon-linked TFs such as STAT1, IRF7, but also GATA6, which is associated with mesenchymal-like states in various cancers [34,35]. The mesenchymal-like TNBC cluster had activation of EMT-related TFs, such as TWIST1 and ZEB1, and the hypoxia-response TNBC cluster had activation of TFs associated with redox homeostasis, like CEBPG, MAFG and KLF5 [36–38]. Transcription factors of note that are active in LP-like and stem-like TNBC clusters include the oncogene MYC, the stress-related TF ATF4, associated with aggressive TNBC [39], and members of the basic helix-loop-helix family, including HEY1, which has been associated with breast cancer metastasis [40]. We also identified a set of TFs spanning the poorly differentiated and proximal mesenchymal-like, hypoxia-responsive and inflammatory response clusters, including PRRX2 and SOX9, both linked to aggressive forms of numerous cancer types [41–43]. Overall, this analysis more comprehensively describes the cell heterogeneity within TNBC and its microenvironment, and details gene programs which may explain how TNBC arises from luminal progenitor cells and adopts different phenotypes.

### A simplified transcriptome signature predicts patient outcome

To see if the different TNBC and TME cell subtypes could be identified from bulk transcriptomes, we used CIBERSORTx^22^ to deconvolve the TNBC samples (*n* = 177) from the TCGA-BRCA dataset [18]. Close populations were merged (TNBC.LPlike1 and TNBC.LPlike2 to “TNBC.LPlike”; TNBC.MESlike and TNBC.IFN to “TNBC.MESlike-IFN”; TNBC.hypoxia and TNBC.inf to “TNBC.hypoxia-inf”) to reduce the frequency of ambiguous calls. The estimated loads of the single-cell clusters within the sampled tumors were readily identified (**Fig 2a**). We then tested relationships between the relative loads of each cell subtype and patient OS, stratifying tumors into high/low levels of the subtype using the optimal cut-point method of their CIBERSORTx scores, and applying a univariate Cox regression model. We found that a relatively high load of myCAFs predicted poor survival (hazard ratio HR = 4; *p* = 0.015) (**Fig 2b**; **Table S5**). For cell types within the tumor itself, poor survival was significantly associated with high STEMlike loads (HR = 2.64; *p* = 0.019), and nearly significantly associated with higher MES-like/IFN loads (HR = 2.17; *p* = 0.066). Conversely, higher proportions of certain immune cell populations, such as CD8 effector T cells (HR = 0.23; *p* = 0.016) and follicular B cells (HR = 0.28; *p* = 0.038) were associated with improved survival. Since CIBERSORTx scores are not practical to implement, and require full transcriptomes to be applied to new samples, we sought to build simpler gene signatures that could replicate these associations, based only on the top marker genes. In this manner, we were able to derive four ten-gene signatures with significant prognostic value (**Fig 2c**): myCAF-sig10 (*COL11A1, MMP1, LRRC15, WNT2, PLPP4, TDO2, HTRA3, POSTN, ADAMTS2, FAP*; HR = 2.75; *p* = 0.014), STEMlike-sig10 (*CHST1, BEST4, SLURP1, ALDH1A1, RBP4, HEY1, DLX1, USH1C, CNTD2, CALML3*; HR = 2.49; *p* = 0.021), and MESlike-sig10 (*RGS2, SPARC, KLK5, CDKN1C, A2M, ATCG2, TIMP3, COL6A2, ACTA2, FBLN2*; HR = NA; *p* = 0.04) for reduced survival, and CD8eff-sig10 (*GZMA, NKG7, CST7, CCL4, TRAC, CD8A, TRBC2, CD3G, GIMAP7*; HR = 0.26; *p* = 0.007) for increased survival. Notably, the ten-gene signature was able to determine a significant association between MESlike cells and poor prognosis, unlike the absolute CIBERSORTx score.

**Figure 2.**
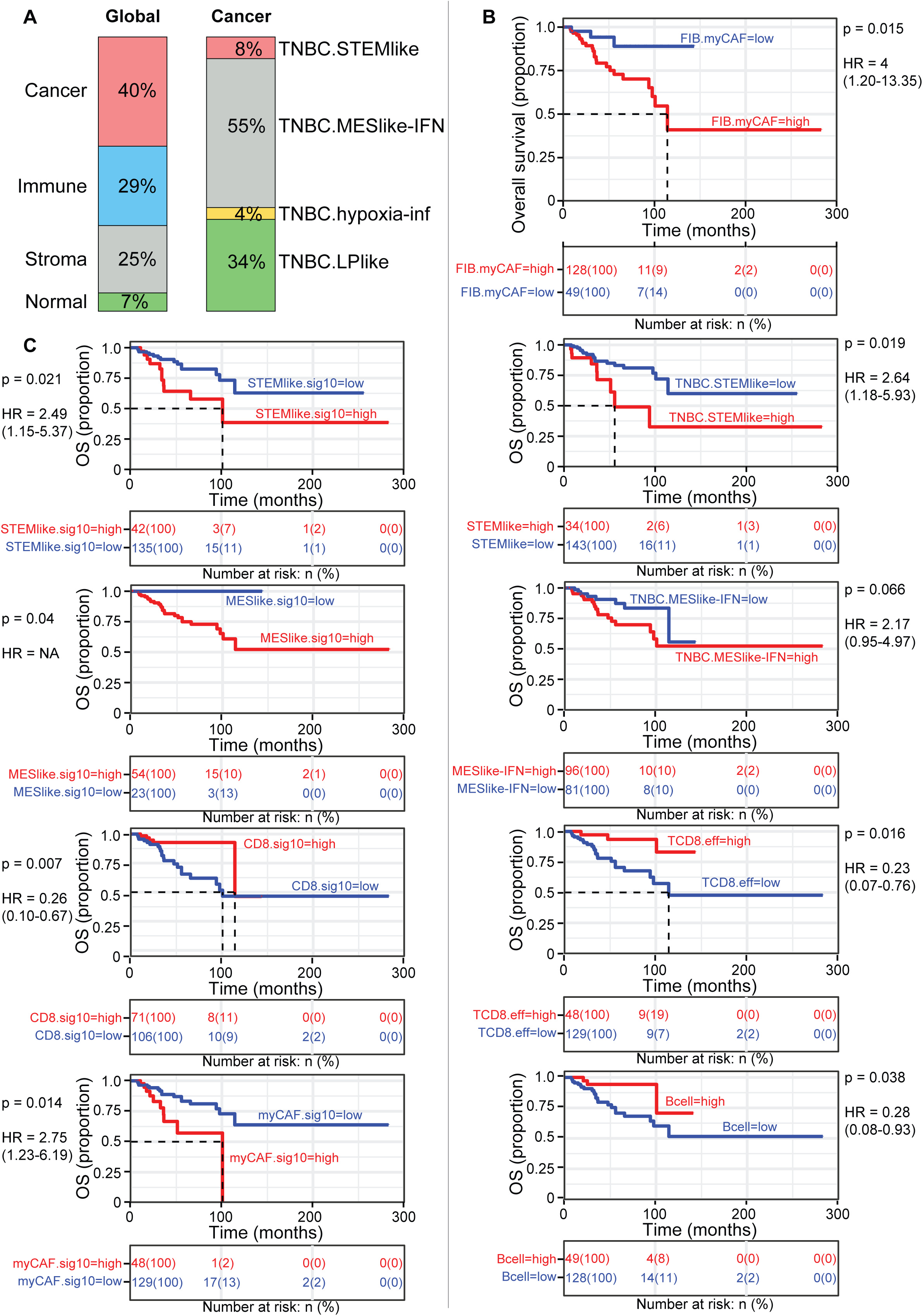
Cell subtypes and their gene signatures associated with overall survival in TNBC. **a**) Average relative proportion of different cell types within TNBC samples from the TGCA-BRCA bulk RNA-seq dataset^18^, inferred by CIBERSORTx deconvolution from the single-cell expression signature clusters identified in analysis of the data from ref 12. **b**) Kaplan-Meier curves for survival analysis of TCGA-BRCA TNBC samples, stratified by high or low burden of denoted cell types, as inferred by CIBERSORTx deconvolution using the optimal cut-off value method. **c**) Kaplan-Meier curves for survival analysis of TCGA-BRCA TNBC samples, stratified by high or low simplified ten-gene signatures from various cell types using the optimal cut-off value method.

### Altered cell type distributions in metastasis

To see how the distributions of different TNBC and TME cell types may differ in metastasis, we applied a similar deconvolution method to bulk transcriptome data from matched primary tumors (*n* = 18) and metastatic sites (*n* = 34) from brain, liver, lung and lymph node [19]. Globally, metastatic samples had a higher proportion of cancer cells and a lower proportion of immune cells, with a comparable proportion of stroma, although there was some variability between patients, as well as between secondary metastatic sites within the same patient (**Fig 3a**; **Fig S4**). Notably, when focusing on the TNBC compartment, the proportions of stem-like and mesenchymal-like cells were also increased at metastatic sites (**Fig 3b**). In line with this, metastases had significantly higher average absolute CIBERSORTx scores for TNBC.STEMlike (log-fold change logFC = 3.65; *p* = 0.002) and TNBC.MESlike-IFN (logFC = 1.38; *p* = 0.0004) subtypes, but also an increase in hypoxic-like CAFs (logFC = 2.82; *p* = 0.03), as well as *IL7R*+ CD4 T cells (logFC = 1.76; *p* = 0.003) (**Fig 3c**; **Table S6**). Metastatic samples had significantly lower CIBERSORTx scores for various lymphocyte subtypes, consistent with the presumed protective role of tumor-infiltrating lymphocytes in TNBC [44]. Overall, these results support the predominant view that stem-like and mesenchymal tumor cells, as well as hypoxic CAFs within the TME, are responsible for malignancy, suggesting that they may even be directly involved in metastatic colonization.

**Figure 3.**
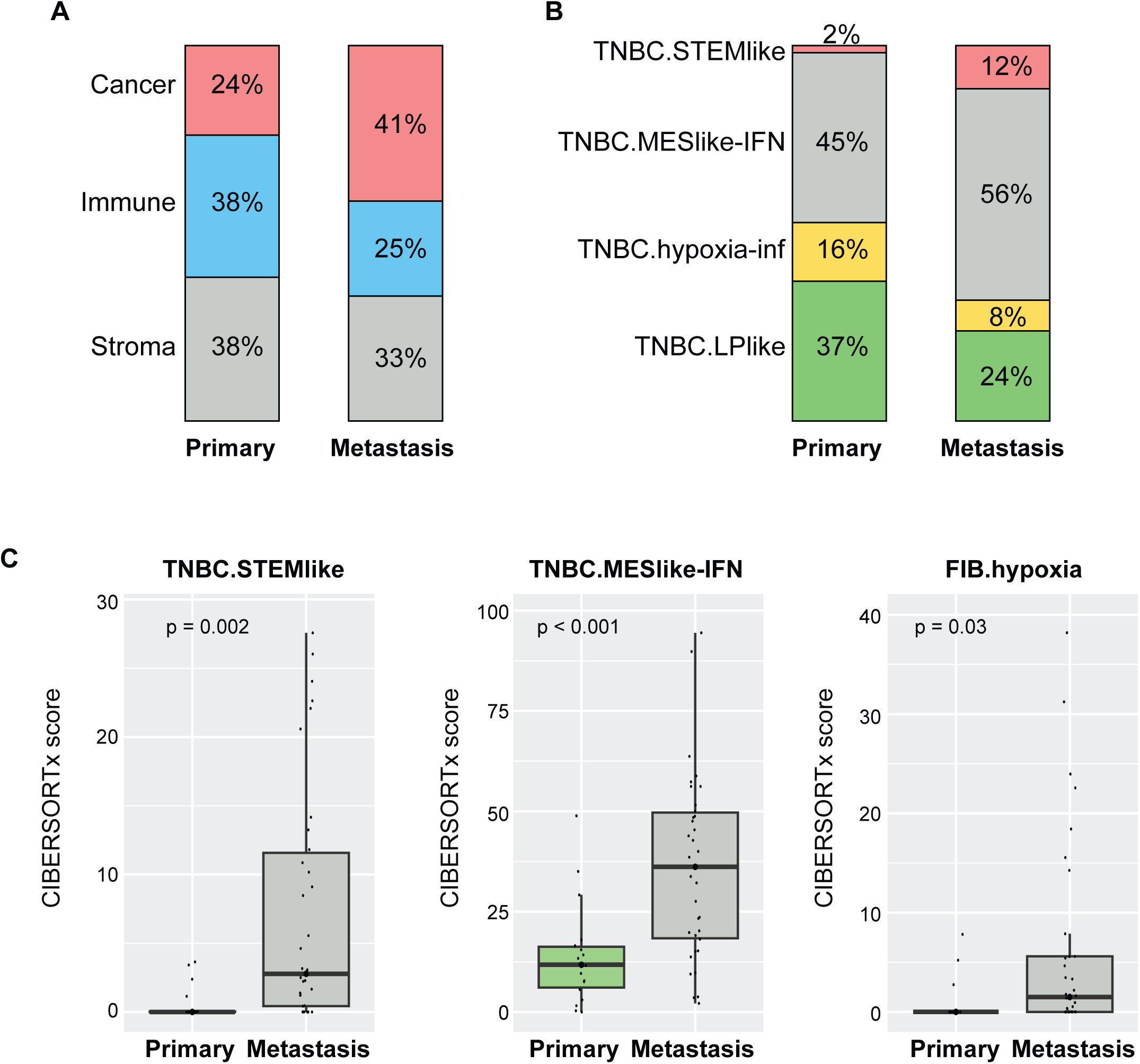
Cell subtypes associated with metastasis in TNBC. **a**) Average relative proportions of different major cell types within primary and metastatic TNBC samples (from ref 19), inferred from bulk RNA-seq data by CIBERSORTx deconvolution from the single-cell expression signature clusters identified in analysis of the data from ref 12. **b**) Average relative proportions of TNBC epithelial cell types within primary and metastatic TNBC samples (from ref 19), inferred from bulk RNA-seq data by CIBERSORTx deconvolution from the single-cell expression signature clusters identified in analysis of the data from ref 12. **c**) Box plots showing distributions of CIBERSORTx scores for denoted cell types, comparing primary and metastatic tumor samples.

### A simplified transcriptome signature predicts therapy response

Since estimating the proportion of specific cell types from transcriptomic deconvolution could predict patient OS and distinguish metastatic from primary tumors, we reasoned that it could also predict patient responses to different therapies. To assess patient responses to neoadjuvant chemotherapy, we used transcriptomes of pre-and post-treatment TNBC samples from the SCANDARE biobank, comprising 62 chemosensitive (49 pre-treatment, 13 post-treatment) and 52 chemoresistant (29 pre-treatment, 23 post-treatment) samples [20]. As expected, cancer cell proportions were greatly reduced in post-treatment chemosensitive samples and not chemoresistant ones (**Fig 4a**). Interestingly, the residual tumor cells in post-treatment chemosensitive samples were slightly enriched in STEMlike and MESlike-IFN cells (**Fig 4b**), suggesting that these residual cancer cells might be TNBC phenotypes most resistant to chemotherapy. We then compared CIBERSORTx scores in pre-treatment chemosensitive and chemoresistant samples to determine if we could find a signature distinguishing capacity to respond to neoadjuvant chemotherapy. In line with initial findings of the SCANDARE study [20], stem-like and mesenchymal-like TNBC tumor cells, as well as myCAFs, were not noticeably enriched in pre-treatment chemoresistant samples (**Fig 4b,c**; **Table S7**). Surprisingly, adipogenic CAFs arose as a cell type significantly enriched in chemoresistant samples (logFC = 1.5; *p* = 0.006). As before, we were able to derive a simpler ten-gene signature for adipogenic CAFs (*APOD, IGF1, CXCL14, ITM2A, CFD, SELENOP, PTGDS, OGN, MGP, NFIA*) that is also significantly enriched in chemoresistant pre-treatment samples (logFC = 0.77; *p* = 0.01) (**Fig 4d**; **Fig S5a**). We note that a similar type of CAF was found to be upregulated in pancreatic cancer samples resistant to chemotherapy [45].

**Figure 4.**
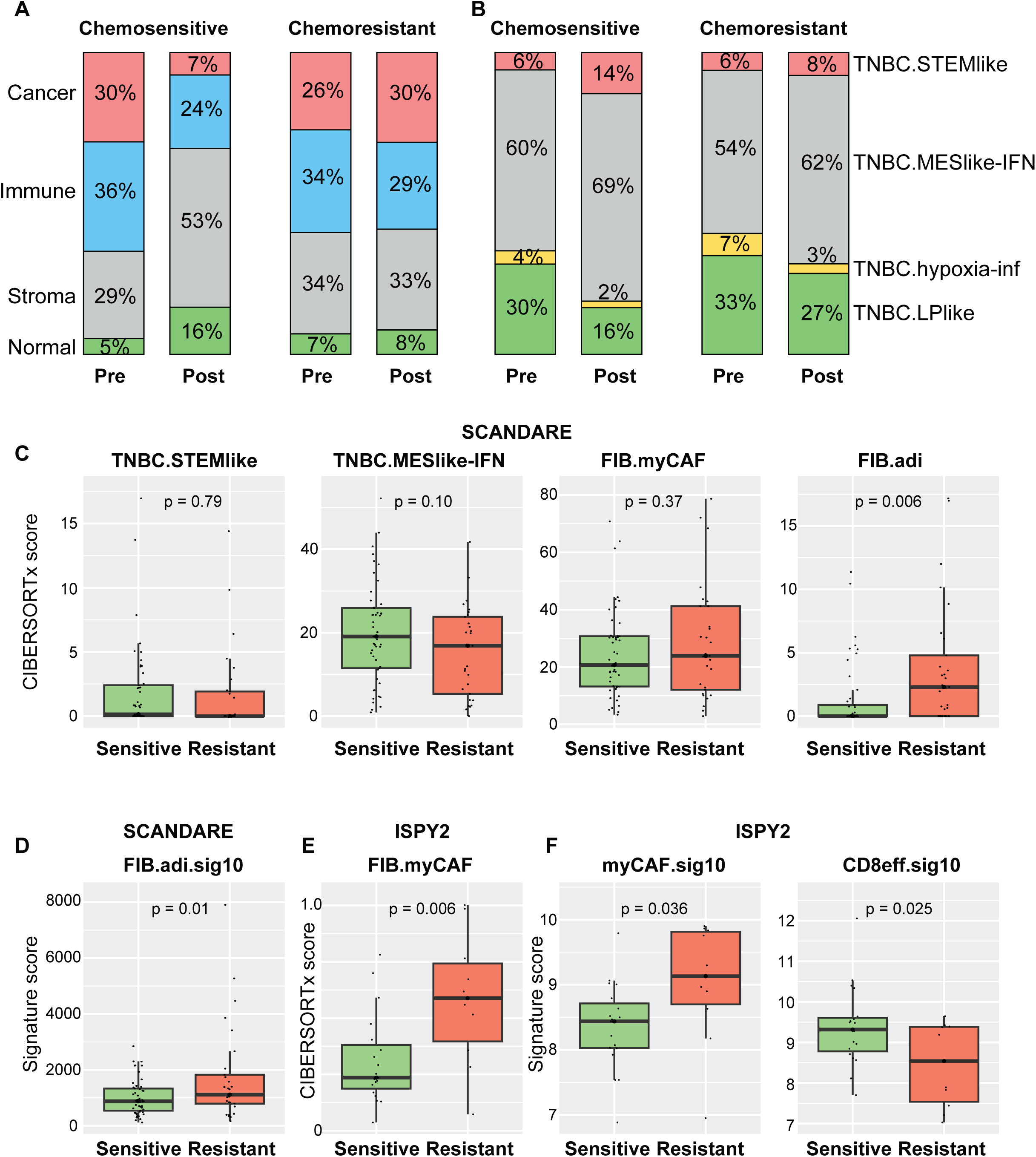
Cell subtypes and gene signatures predicting response to different therapy strategies. **a**) Average relative proportions of different major cell types within TNBC samples determined to be either sensitive or resistant to neoadjuvant chemotherapy, taken before or after treatment, inferred from bulk RNA-seq data in the SCANDARE study^20^ by CIBERSORTx deconvolution from the single-cell expression signature clusters identified in analysis of the data from ref 12. **b**) Average relative proportions of TNBC epithelial cell types from samples determined to be either sensitive or resistant to neoadjuvant chemotherapy, taken before or after treatment, inferred from bulk RNA-seq data in the SCANDARE study^20^ by CIBERSORTx deconvolution from the single-cell expression signature clusters identified in analysis of the data from ref 12. **c**) Box plots showing distributions of CIBERSORTx scores for denoted cell types, comparing pre-treatment tumor samples subsequently found to be sensitive or resistant to neoadjuvant chemotherapy^20^. **d**) Box plot showing distributions of simplified ten-gene signature scores predicting adipogenic CAF burden, comparing pre-treatment tumor samples subsequently found to be sensitive or resistant to neoadjuvant chemotherapy^20^. **e**) Box plot showing distributions of CIBERSORTx scores for predicted myCAF burden, derived from deconvolution of microarray expression analysis on the ISPY2 study^21^, comparing pre-treatment tumor samples subsequently found to be sensitive or resistant to chemo-immunotherapy with paclitaxel and pembrolizumab. **f**) Box plots showing distributions of simplified ten-gene signature scores predicting burdens of denoted cell subtypes, derived from deconvolution of microarray expression analysis on the ISPY2 study^21^, comparing pre-treatment tumor samples subsequently found to be sensitive or resistant to chemo-immunotherapy with paclitaxel and pembrolizumab.

To assess cell types that may influence responses to chemo-immunotherapy, which were not included in the SCANDARE biobank, we analyzed microarray-derived transcriptome data from the I-SPY2 clinical trial [21], focusing on the pembrolizumab-containing neoadjuvant regimen, which is the current standard of care for early TNBC by the KEYNOTE-522 study [46]. Comparing CIBERSORTx scores for responder (*n* = 19) and non-responder (*n* = 10) cases, we found that responders had higher scores for some immune cell types, as may be expected for response to immune checkpoint inhibitors (**Table S8**). Most notably, non-responders had a significantly increased score for myCAFs (logFC = 0.91; *p* = 0.006), but not for adipogenic CAFs (logFC =-0.57; *p* = 0.65), stem-like (logFC = 0.6; *p* = 0.24) or mesenchymal-like (logFC =-0.07; *p* = 0.79) tumor cells (**Fig 4e**; **Fig S5b**). Furthermore, the same ten-gene signatures which we used to associate myCAF or CD8 effector T cell load with reduced or improved overall survival, respectively, also demonstrated significant differences between responders and non-responders (myCAFs *p* = 0.034; CD8 effector T cells *p* = 0.025) (**Fig 4f**). Overall, this suggests that prevalence of CAFs can confer resistance of TNBC to systemic therapy, with adipogenic CAFs particularly impeding neoadjuvant chemotherapy and myCAFs particularly impeding immunotherapy. Importantly, these differential sensitivities could potentially be identified in the clinic from a simple transcriptional signature.

## DISCUSSION

Our single-cell analysis of TNBC tumors profiled 43 populations, including 6 TNBC-specific epithelial cell phenotypes, and the stromal and immune cells in the associated TME. In line with previous studies [30–32], we find luminal progenitors as the most likely origin of TNBC. Through differential gene expression analysis we identified downregulation of *OVOL2* and activation of stress response pathways as possible drivers of pre-neoplastic state, with subsequent activation of known oncogenic drivers for multiple cancer types, such as the NOTCH and WNT pathways, and transcription factors of the AP-1, SOX and SIX families, linked to full transformation. Recapitulating the findings of many previous studies [11,47,48], poor survival and/or increased invasiveness was found to be associated with a higher load of stem-cell like and mesenchymal tumor cells, as well as increased myCAF or hypoxic CAF load in the TME, whereas better prognosis is correlated with a higher load of tumor-infiltrating lymphocytes. Original bulk transcriptomic analyses of TNBCs attempted to classify tumors into certain types, such as “basal” or “luminal androgen receptor” [9]. Some clinical data suggest that anti-androgens could be a viable therapy for TNBC cases characterized in this way as “LAR” based on IHC assessment [49]. However, such classifications have had limited prognostic value, particularly as cells with the expression signature of all of these types were found to be present within most tumors when studied at the single-cell level [29]. Instead, deconvolution methods to estimate relative proportions of the aggressive or protective cell types within the tumor have provided clinically relevant statistical metrics [50, 51]. We have extended this effort further to identify simpler ten-gene signatures that provide similar predictions of TNBC OS without the need for single-cell or even bulk transcriptome data collection of new clinical cases.

Curiously, although residual tumors after chemotherapy contain on average greater proportions of stem-like and mesenchymal TNBC than pre-treatment tumors, we found no significant association between these cancer-intrinsic cell types and response to therapy. As noted in the original study [20], the composition of the stroma appeared more influential in determining therapy response. With the small sample sizes available in the analyzed dataset, we could not rule out potential differences between different metastatic sites. Our deconvolution results indicate that CAFs are a key component of the TME across samples. In particular, myCAFs are the population giving the highest hazard ratio, and are capable of stratifying patients that do not respond to chemo-immunotherapy. Interestingly, myCAFs were only slightly and non-significantly enriched in tumors resistant to neoadjuvant chemotherapy; instead, adipogenic CAFs seem to play a greater role in resistance to this treatment strategy. These latter results highlight how characterization of the TME can inform not just the gravity of the disease, but could potentially inform the most effective treatment option, the main goal of personalized medicine.

Our analysis has a major limitation: a limited sample size for clinical outcome measurements and the absence of a validation cohort. Despite this, we were able to reduce the predictive signatures down to a small subset of genes (ten genes each, for five key cell types). Further studies of larger cohorts, and prospective validation, will be required to formally assess the power of this approach, particularly when combining with characteristics already used in the clinic, including biomarkers such as TILs or PD-L1. Despite demonstrating significant improvements in event-free and overall survival, the KEYNOTE-522 neoadjuvant regimen currently lacks a reliable biomarker predictive of response or resistance; our relatively simple estimate of myCAF burden warrants further investigation as potentially such a marker. The second limitation is the absence of consideration of spatial distribution [52, 53] and cell-cell interactions between identified tumoral subtypes and myCAFs [54], which have been demonstrated to have prognostic value. However, this study raises the possibility that personalized treatment strategies for breast cancer could be informed in the clinic from much more basic and less costly qRT-PCR assays, without the need for costly single-cell (or even bulk) or spatial transcriptomics/proteomics approaches. In principle, use of these stromal signatures could be expanded to other subtypes of breast cancer and other cancers treated using chemotherapy and immune checkpoint inhibitors.

## ABBREVIATIONS

CAF: Cancer-associated fibroblast
CSC: Cancer stem cell
EMT: Epithelial-mesenchymal transition
ER+/PR+: Estrogen receptor/progesterone receptor-positive
HR: Hazard ratio
IFN: Interferon
logFC: Log-fold change
LP: Luminal progenitor
ML: Mature lumen
myCAF: Myofibroblastic cancer-associated fibroblast
TAM: Tumor-associated macrophage
TME: Tumor microenvironment
TNBC: Triple negative breast cancer
UMAP: Uniform manifold approximation and projection

## DECLARATIONS

### Ethics approval and consent to participate

Not applicable

### Consent for publication

Not applicable

### Availability of data and materials

The single-cell RNA-sequencing data are available at Gene Expression Omnibus (GSE161529). The bulk RNA-sequencing data from the TCGA-BRCA collection are available from the GDC data portal (https://portal.gdc.cancer.gov/projects/TCGA-BRCA). The bulk RNA-sequencing data for the metastatic study are available at Gene Expression Omnibus (GSE209998). The bulk RNA-sequencing data from the SCANDARE study (NCR03017573) are available from EGA (EGAS50000000970). The microarray data from the I-SPY2 clinical trial are available at the Gene Expression Omnibus (GSE194040). All meta-analyses are available in the Supplementary Tables of this manuscript.

### Contributions

G.D. and T.S. conceptualized the project. G.D. performed all the analyses, with input from V.D. for the survival analyses. G.D. provided data visualization. G.D. and T.S. wrote the manuscript, with critical review from V.D. All authors read and approved the final version of the manuscript.

