## Supplementary figures and images for "Single-cell analyses reveal a simple multi-gene transcriptomic signature with predictive power in prognosis and therapy effectiveness in triple-negative breast cancer"

### Supplemental Fig 1

**Davidson et al., Figure S1**

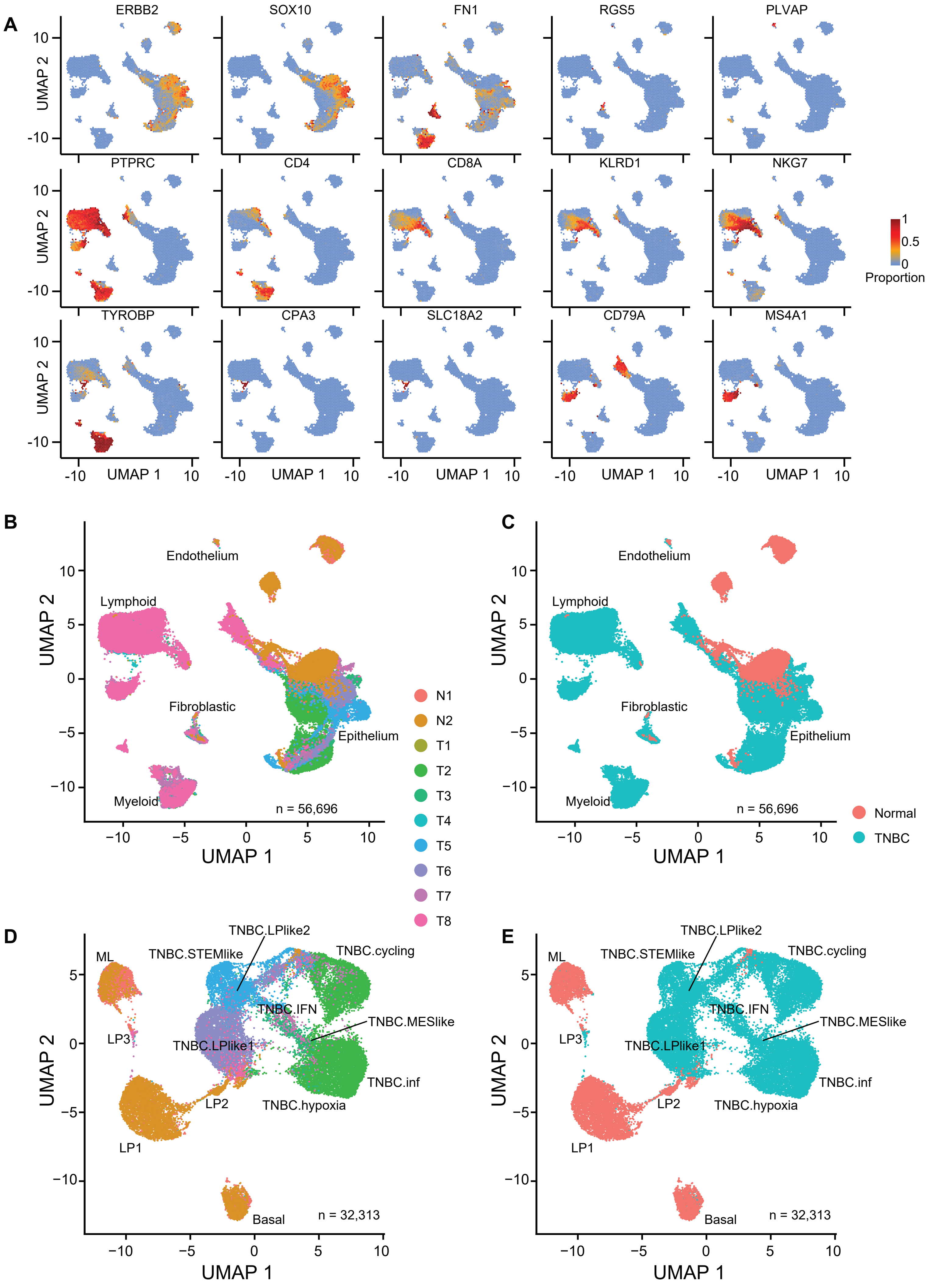

### Supplemental Fig 2

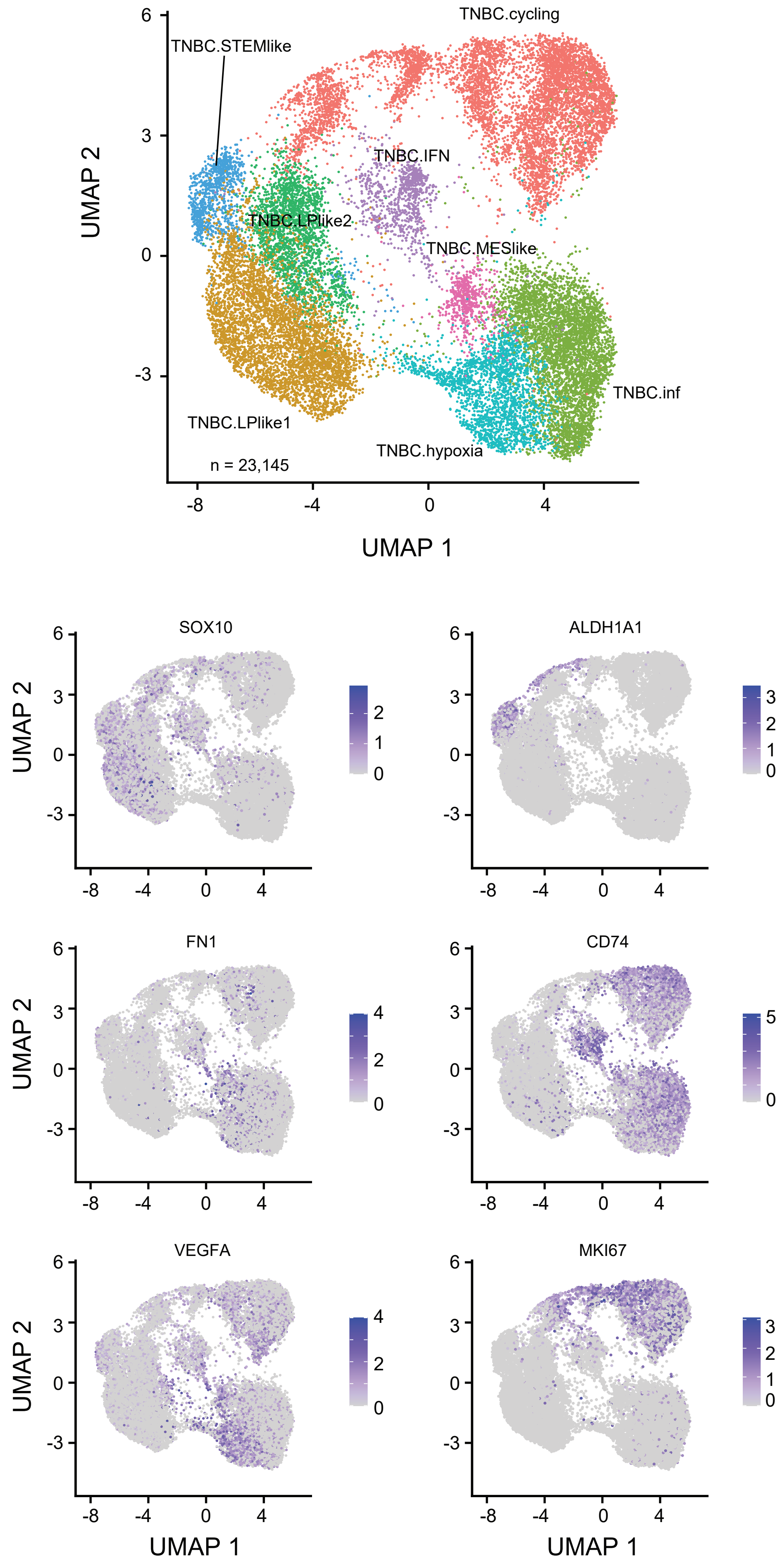

### Supplemental Fig 3

**Davidson et al., Figure S3**

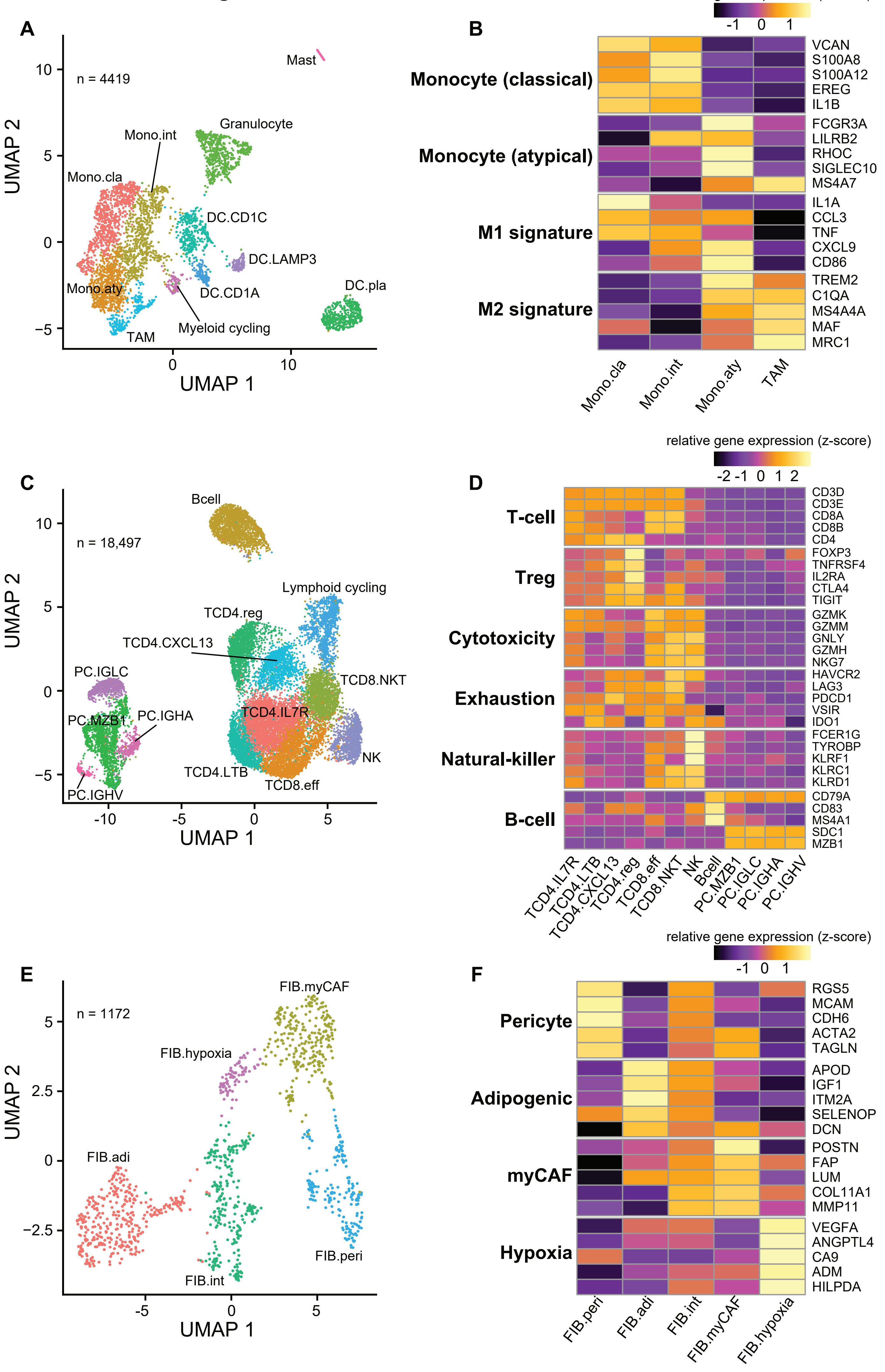

### Supplemental Fig 4

Davidson et al., Figure S4

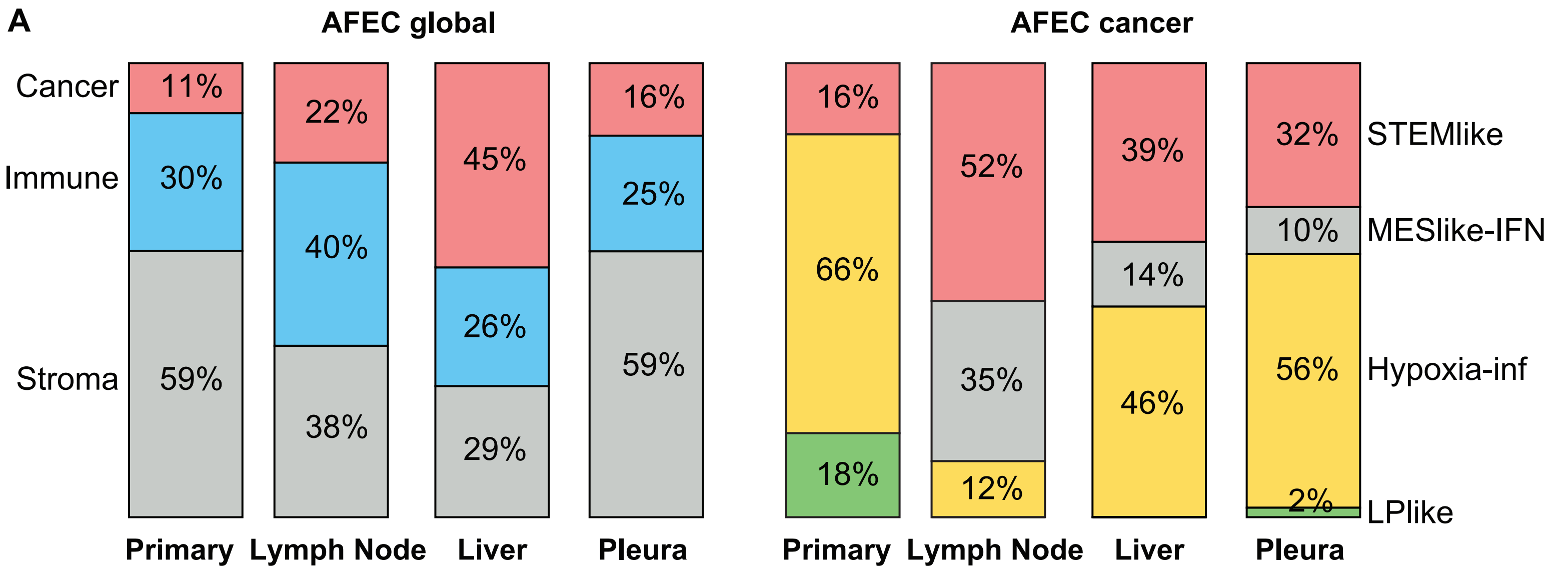


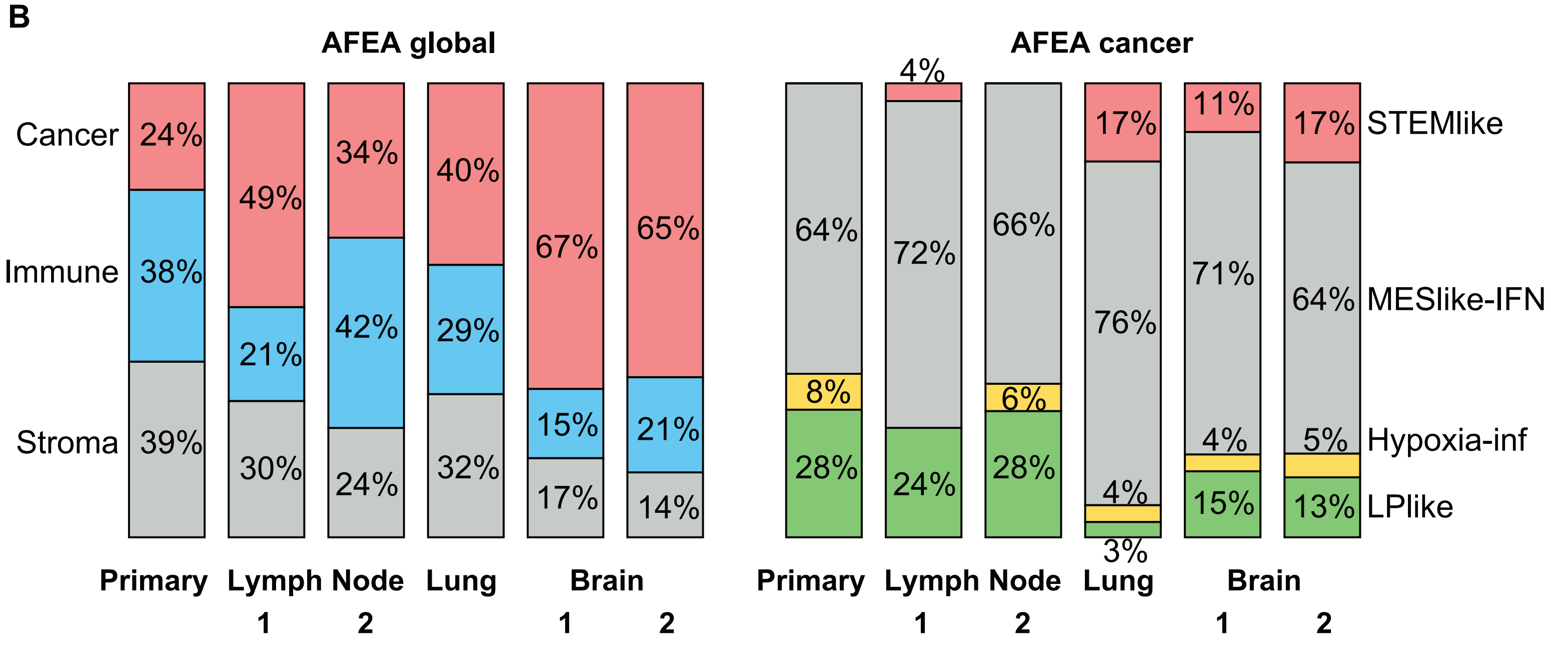


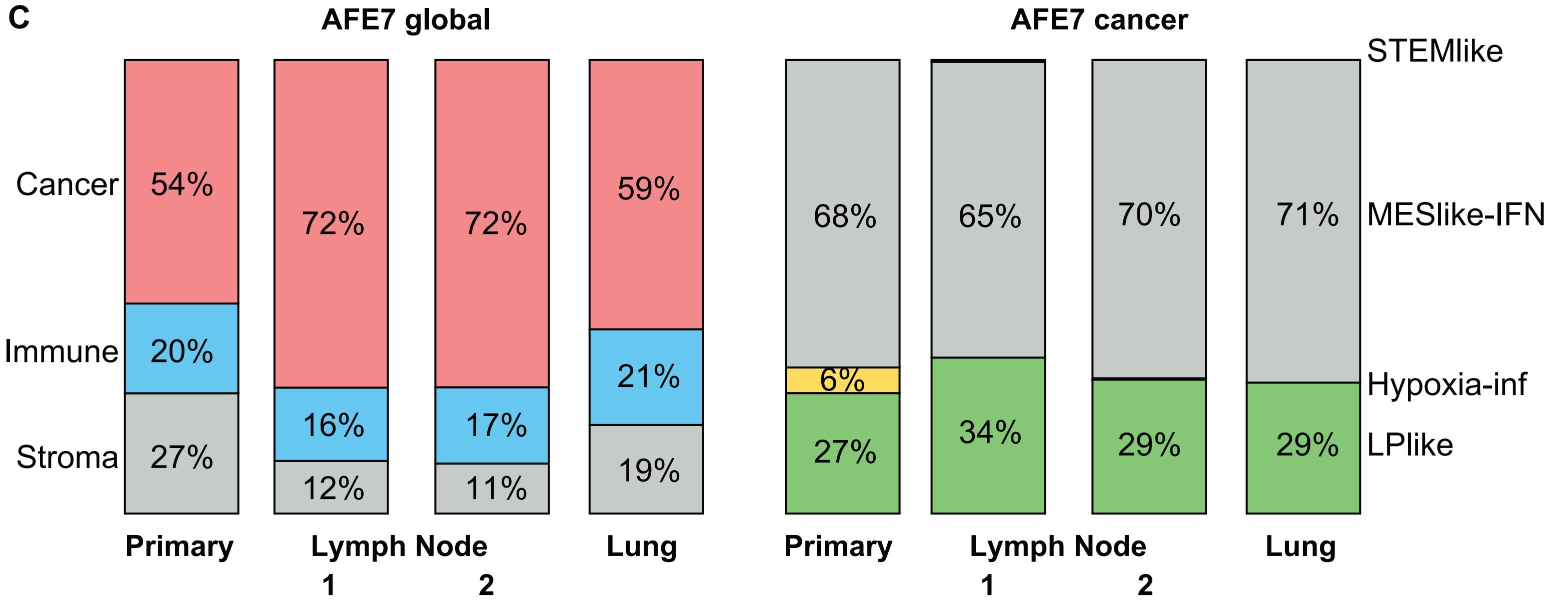

### Supplemental Fig 5

Davidson et al., Figure S5

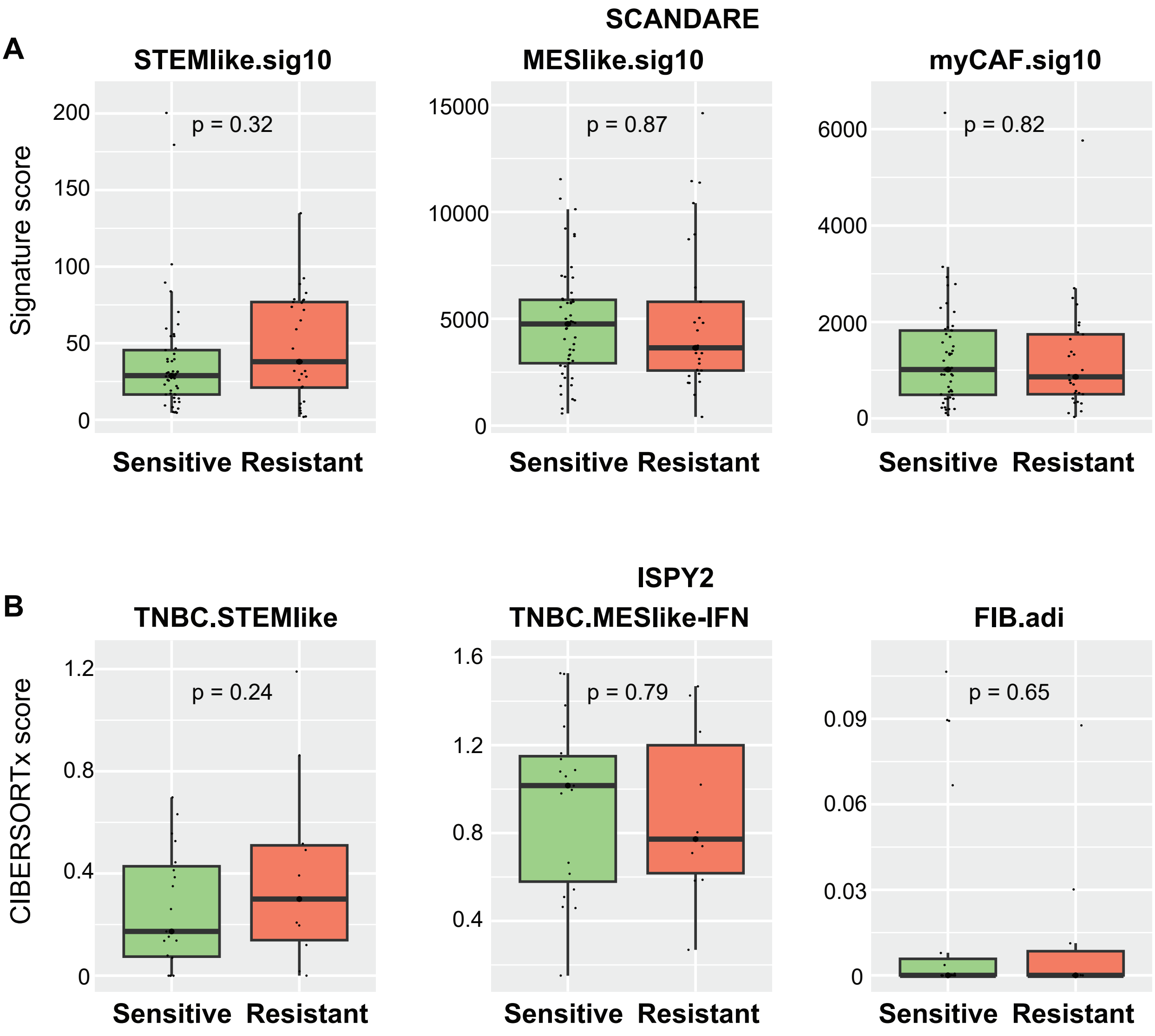
