## Supplemental Text and Table 1 for "Single-cell analyses reveal a simple multi-gene transcriptomic signature with predictive power in prognosis and therapy effectiveness in triple-negative breast cancer"

**SUPPLEMENTARY DATA**

**SUPPLEMENTARY FIGURES**

**Fig S1. Batch representation of TNBC cell subtypes. a**) UMAP projections and clustering of single-cell transcriptomic data (from ref 12), with indications of the proportions of cells expressing denoted marker genes. **b**) UMAP projections and clustering of single-cell transcriptomic data (from ref 12), color-coded to denote the sample origins of each cell (N, normal sample; T, tumor sample). **c**) As **b**), but color-coded according to normal or TNBC origin of each cell. **d**, **e**) As **b, c**), but for re-clustered epithelial cells.

**Fig S2. Gene markers of epithelial cell subtypes within TNBC and normal breast.** UMAP projections and clustering of EPCAM^+^ (epithelial) portion of cells from single-cell transcriptomic data (ref 12). Below, the same UMAP projections are shown, color-coded with logFC for expression of denoted marker genes.

**Fig S3. Diversity of myeloid, lymphoid and fibroblast cell types within TNBC TME.** **a**) UMAP projections and re-clustering of myeloid portion of cells from single-cell transcriptomic data (ref 12). **b**) Heat map showing the relative expression levels of given marker genes within the different myeloid clusters, identified in **a**). **c,d**) As **a,b**) for the lymphoid compartment. **e,f**) As **a,b**) for the fibroblast compartment.

**Fig S4. Heterogeneity of primary and metastatic TNBC samples.** **a**) Relative proportions of different major cell types (left) and TNBC epithelial cell types (right) within primary and different metastatic site tumor samples from individual patient AFEC (ref 19), inferred from bulk RNA-seq data by CIBERSORTx deconvolution from the single-cell expression signature clusters identified in analysis from the data of ref 12. **b**) As **a**) for patient AFEA. **c**) As **a**) for patient AFE7.

**Fig S5. Poor predictive value of TNBC epithelial subtype cell burden to therapy response. a**) Box plots showing distributions of simplified ten-signatures predicting denoted cell type burdens, comparing pre-treatment tumor samples subsequently found to be sensitive or resistant to neoadjuvant chemotherapy (ref 20). **b**) Box plots showing distributions of CIBERSORTx scores for denoted cell types, comparing pre-treatment tumor samples subsequently found to be sensitive or resistant to immunotherapy (ref 21).

**SUPPLEMENTARY TABLES**

**Table S1. Cell types within TNBC tumor and TME.** Descriptions of the different cell types identified in single-cell transcriptome analysis of TNBC samples (ref 12).

| **Cluster name** | **Cell type (broad)** | **Description** |
| --- | --- | --- |
| Basal | Normal breast epithelium | Breast basal cells |
| LP1 | Normal breast epithelium | Breast luminal progenitors |
| LP2 | Normal breast epithelium | Breast luminal progenitors, preneoplastic |
| LP3 | Normal breast epithelium | Breast luminal progenitors, differentiating into mature luminal cells |
| ML | Normal breast epithelium | Breast mature luminal cells |
| TNBC.LPlike1 | Cancer cells | TNBC cells, LP-like phenotype, closest identity to cell of origin |
| TNBC.LPlike2 | Cancer cells | TNBC cells, LP-like, dedifferentiating towards a stem-like phenotype |
| TNBC.STEMlike | Cancer cells | TNBC cells, stem-like phenotype |
| TNBC.IFN | Cancer cells | TNBC cells, interferon-response phenotype |
| TNBC.inf | Cancer cells | TNBC cells, inflamed phenotype |
| TNBC.hypoxia | Cancer cells | TNBC cells, hypoxia-response phenotype |
| TNBC.MESlike | Cancer cells | TNBC cells, mesenchymal-like phenotype |
| TNBC.cycling | Cancer cells | TNBC cells, going through cell cycle |
| FIB.peri | Fibroblasts | Pericytes |
| FIB.adi | Fibroblasts | Fibroblasts, adipogenic CAF precursors derived from adipocytes |
| FIB.int | Fibroblasts | Fibroblasts, intermediate between adipogenic precursors and CAFs |
| FIB.hypoxia | Fibroblasts | Fibroblasts, hypoxia-response phenotype |
| FIB.myCAF | Fibroblasts | Cancer-associated fibroblasts, myofibroblastic type |
| Mono.cla | Myeloid cells | Monocytes, classical phenotype |
| Mono.int | Myeloid cells | Monocytes, intermediate phenotype |
| Mono.aty | Myeloid cells | Monocytes, atypical (non-classical) phenotype |
| TAM | Myeloid cells | Tumor-associated macrophages, M2 phenotype |
| DC.CD1C | Myeloid cells | Dendritic cells, conventional phenotype |
| DC.CD1A | Myeloid cells | Dendritic cells, IL-12-producing phenotype |
| DC.LAMP3 | Myeloid cells | Dendritic cells, immuno-regulatory phenotype |
| DC.pla | Myeloid cells | Dendritic cells, plasmacytoid type |
| Granulocyte | Myeloid cells | Granulocytes |
| Myeloid.cycling | Myeloid cells | Myeloid cells, going through cell cycle |
| Mast | Myeloid cells | Mast cells |
| TCD4.IL7R | Lymphoid cells | CD4+ T cells, memory/naïve with high expression of IL7R |
| TCD4.LTB | Lymphoid cells | CD4+ T cells, memory/naïve with high expression of LTB |
| TCD4.CXCL13 | Lymphoid cells | CD4+ T cells, high expression of CXCL13 associated with tertiary lymphoid structures |
| TCD4.reg | Lymphoid cells | CD4+ T cells, Treg phenotype |
| TCD8.eff | Lymphoid cells | CD8+ T cells, cytotoxic effector phenotype |
| TCD8.NKT | Lymphoid cells | CD8+ NK-like T cells, cytotoxic with signs of exhaustion (HAVCR2, LAG3) |
| NK | Lymphoid cells | Natural killer cells |
| Bcell | Lymphoid cells | B cells, follicular |
| PC.MZB1 | Lymphoid cells | Plasma cells |
| PC.IGLC | Lymphoid cells | Plasma cells, high expression of IGLC immunoglobulins |
| PC.IGHA | Lymphoid cells | Plasma cells, high expression of IGHA immunoglobulins |
| PC.IGHV | Lymphoid cells | Plasma cells, high expression of IGHV immunoglobulins |
| Lymphoid.cycling | Lymphoid cells | Lymphoid cells, going through cell cycle |
| ED | Endothelium | Endothelial cells |

**Table S2. Marker genes of TNBC cell subtypes.** Lists of genes whose expression is significantly associated with a particular TNBC cell type cluster, with average log2-fold change of expression level and adjusted p-values. Accompanying Excel sheet.

**Table S3. Genes upregulated in LP2 compared to LP1.** List of upregulated genes on LP1-to-LP2 transition, with associated average log2-fold change of expression and adjusted p-values. Accompanying Excel sheet.

**Table S4. Genes upregulated in TNBC.LPlike1 compared to LP2.** List of upregulated genes on LP2-to-TNBC.LPlike1 transition, with associated average log2-fold change of expression and adjusted p-values. Accompanying Excel sheet.

**Table S5. TNBC cell subtype links to overall survival.** Hazard ratios for each TNBC cell subtype after their deconvolution from TCGA-BRCA cohort data (ref 18). Accompanying Excel sheet.

**Table S6. Different composition of TNBC cell subtypes in primary and metastatic tumors.** Log2-fold change of expression and p-values for differential burdens of TNBC cell subtypes, comparing primary and metastatic samples (ref. 19). Accompanying Excel sheet.

**Table S7. Different composition of TNBC cell subtypes in tumors sensitive or resistant to neoadjuvant chemotherapy.** Log2-fold change of expression and p-values for differential burdens of TNBC cell subtypes, comparing samples sensitive and resistant to neoadjuvant chemotherapy in the SCANDARE study (ref 20). Accompanying Excel sheet.

**Table S8. Different composition of TNBC cell subtypes in tumors sensitive or resistant to immunotherapy.** Log2-fold change of expression and p-values for differential burdens of TNBC cell subtypes, comparing samples sensitive and resistant to immunotherapy (pembrolizumab) in the ISPY2 study (ref 21). Accompanying Excel sheet.
